# Bcl-3 suppresses Th9 differentiation by regulating glutamine utilization

**DOI:** 10.1101/2021.07.06.451316

**Authors:** Wanhu Tang, Hongshan Wang, Philip M. Murphy, Ulrich Siebenlist

**Affiliations:** Laboratory of Molecular Immunology, National Institute of Allergy and Infectious Diseases, National Institutes of Health, Bethesda, MD 20892, USA

**Keywords:** NF-κB, Bcl-3, Th9

## Abstract

Bcl-3 is an atypical member of the IκB protein family that plays important and diverse roles in both innate and adaptive immunity, including Th17-dependent autoimmunity models in mice. When naïve mouse splenic CD4^+^ T cells were cultured under Th17 conditions *in vitro*, we unexpectedly found that the most highly differentially expressed gene between wild type and Bcl-3-deficient (KO) Th17 cells encoded the cytokine IL-9. We therefore investigated the role of Bcl-3 in Th9 cell differentiation. When naïve CD4^+^ T cells were cultured under Th9-polarizing conditions *in vitro*, the extent of Th9 differentiation observed in wild type cells was increased in Bcl-3 KO cells and conversely was decreased in cells overexpressing Bcl-3. The suppressive effect of Bcl-3 on Th9 differentiation was cell-autonomous, and NF-κB inhibitors abolished increased Th9 differentiation in Bcl-3 KO cells. Consistent with this, in the Th9 transfer model of OVA-induced allergic airway inflammation, mice receiving Bcl-3 KO cells had greater immune cell infiltration in the lung than mice receiving wild type cells.

Mechanistically, unsupervised transcriptomic analysis revealed differentially expressed genes in KO cells, including the glutamine transporter *Slc1a5*, which was downregulated. The functional significance of this was suggested by the ability of increasing concentrations of glutamine in the media to reduce the difference in Th9 differentiation between WT and KO cells. Our results suggest a novel role for Bcl-3 as a negative regulator of Th9 differentiation, in part by limiting glutamine accessibility through downregulation of *Slc1a5*.

## Introduction

Bcl-3 plays important roles in various physiological processes (1-4). It was first cloned as an oncogene involved in the genomic translocation responsible for a form of B cell lymphoma (Bcl). Structural and biochemical studies identified Bcl-3 as an atypical IκB protein, containing both an ankyrin domain and a transactivation domain. Classical IκBs exert their inhibitory function by binding to and retaining the NF-κB transcription factor complex in the cytoplasm; however, Bcl-3 can enter the nucleus and modulate the activity of NF-κB bound to DNA. Bcl-3 mainly binds to NF-κB1 p50 or NF-κB2 p52 homodimers, and it may either promote or inhibit NF-κB-regulated gene expression depending on the context (5-7). Bcl-3 is critical in both innate and adaptive immunity and is expressed in multiple cell types, including B and T cells, DCs and stromal cells (8-13). Previous studies using Bcl-3-deficient mice showed that Bcl-3 was involved in different disease models, including diabetes, rheumatoid arthritis, colitis and experimental autoimmune encephalitis (EAE) (14-18). In the clinic, Bcl-3 has been linked to the pathogenesis of inflammatory bowel disease (IBD) and certain cancers (16, 17, 19-21).

Th9 cells are IL-9-secreting T helper cells that can be generated *in vitro* by culturing CD4^+^ T cells with the cytokines IL-4 and TGFβ (22). They have been demonstrated to be mostly detrimental in different diseases, including airway inflammation, IBD and EAE (23-25) although beneficial roles have also been reported in EAE as well as in certain infectious disease models (25, 26). Th9 cells have also been reported to be more effective than Th1 and Th17 cells in cancer adoptive cell transfer therapy (27). IL-9 acts on distinct types of cells, including B cells, smooth muscle cells, airway epithelial cells and goblet cells (23, 25, 28). Unlike other types of T helper cells, the master transcription factor for Th9 cells has not been clearly defined. BATF, IRF4, IRF8 and PU.1 have all been shown to be involved in Th9 differentiation, but these transcription factors are also expressed in other types of T helper cells (29-32). Previous studies suggested that the coordination of transcription factors downstream of IL-4 and TGFβ signaling may be required for optimal Th9 differentiation (22).

We have previously shown that Bcl-3 is not required for Th1, Th17 and Treg differentiation, but it is essential for Th1 cell plasticity and pathogenicity (8, 33). Here we show that IL-9 is the most highly differentially expressed gene in naïve CD4^+^ wild type versus Bcl-3 knockout T cells cultured under Th17 conditions and therefore we have investigated the role of Bcl-3 in Th9 differentiation *in vitro*, as well as its potential effect on Th9-mediated airway inflammation *in vivo*.

## Materials and Methods

### Mice

Bcl-3-deficient and Bcl-3-overexpressing mice have been previously described (8, 34). Mice were housed in National Institute of Allergy and Infectious Diseases facilities, and all experiments were done with approval of the National Institute of Allergy and Infectious Diseases Animal Care and Use Committee and in accordance with all relevant institutional guidelines.

### T cell differentiation

Naïve CD4^+^ T cells were isolated from spleens of 6- to 8-week old healthy C57BL/6 mice using a Miltenyi Naïve CD4^+^ T cells Isolation Kit (Gaithersburg, MD) and seeded in DMEM at 5 × 10^5^ cells per well of a 96-well plate coated with 1 μg/mL anti-CD3 (145-2C11). 2 μg/mL anti-CD28 (37.51) were added along with the following cytokines and blocking antibodies to establish specific T helper cell differentiation conditions: 100 U/mL IL-2, 20 ng/mL IL-4, 10 ng/mL TGFβ, 10 μg/mL anti-IL-12 (C18.2) and 10 μg/mL anti-IFNγ (XMG1.2) for Th9 conditions; 100 U/mL IL-2, 10 ng/mL IL-12 and 10 μg/mL anti-IL-4 (11B11) for Th1 conditions; 100 U/mL IL-2, 5 ng/mL TGFβ, 10 μg/mL anti-IL-12, 10 μg/mL anti-IFNγ and 10 μg/mL anti-IL-4 for Treg conditions; and 20 ng/mL IL-6, 5 ng/mL TGFβ, 10 μg/mL anti-IL-12, 10 μg/mL anti-IFNγ and 10 μg/mL anti-IL-4 for Th17 conditions. IL-1 (10 ng/mL), IL-21 (50 ng/mL) and IL-23 (20 ng/mL) were added in addition to Th17 reagents for Th17^+^ conditions. T cells were cultured for 3 days before FACS analysis. All cytokines and antibodies noted above were purchased from PeproTech (Rocky Hill, NJ) and BioXCell (West Lebanon, NH), respectively, except IL-6 (R&D, Minneapolis, MN).

### Flow cytometry

Lymph node cell suspensions were filtered with 100 μm filters and stained with surface antibodies for FACS analysis. Perfused lungs were processed as previously described (35). For intracellular staining, cells were first stained with antibodies for surface marker expression, then permeabilized and stained with antibodies for intracellular protein for 1 h using the Foxp3 Staining Buffer Set (Thermo Fisher Scientific, Waltham, MA). Data were collected in a FACSCalibur flow cytometer (BD Biosciences, San Jose, CA) and analyzed using FlowJo software (BD, Franklin Lakes, NJ). Antibodies to the following markers were used: IL-9 (RM9A4, Biolegend, San Diego, CA), CD4 (RM4-5, Thermo Fisher Scientific, Waltham, MA), CD3 (145-2C11, Biolegend, San Diego, CA), TCRβ (H57-597, BD Biosciences, San Jose, CA), TCR Vα2 (B20.1, BD Biosciences, San Jose, CA), Ly5.1 (A20, BD Biosciences, San Jose, CA), Ly5.2 (104, BD Biosciences, San Jose, CA), Foxp3 (FJK-16s, Thermo Fisher Scientific, Waltham, MA), IL-17 (eBio17B7, Thermo Fisher Scientific, Waltham, MA), and IFNγ (XMG1.2, Thermo Fisher Scientific, Waltham, MA).

### Th9 transfer model of OVA-induced allergic airway inflammation

Naïve WT and Bcl-3 KO OTII CD4^+^ T cells (CD45.2) were isolated and cultured under Th9 conditions for 3 days *in vitro*. Differentiated Th9 OTII cells were adoptively transferred into CD45.1 recipients at 5 × 10^6^ cells per mouse intravenously. The recipients were then treated intranasally with OVA (100 μg/mouse/day, Sigma, Burlington, MA) and IL-25 (500 ng/mouse/day, Biolegend, San Diego, CA) every 4 days for 24 days, and sacrificed one day after the last treatment on day 25.

### Immunohistochemistry

For H&E staining, lungs were prepared as described (35) and embedded in paraffin. Sections were cut at 5 μm thickness and stained with H&E. Lung inflammation was graded using a scale of 0 - 6 for perivascular and peri-bronchial cell infiltration.

### Real-time PCR

RNA was isolated using the RNeasy kit (Qiagen, Germantown, MD) according to the manufacturer’s instructions. cDNA was synthesized with Superscript III (Thermo Fisher Scientific, Waltham, MA). Gene expression was quantified with TaqMan real-time PCR primers (Thermo Fisher Scientific, Waltham, MA). The results were normalized against β-actin.

### RNA sequencing

RNA samples were sequenced on HiSeq at NCI Frederic facility using Illumina (San Diego, CA) TruSeq Stranded Total RNA Kit RS-122-2201 and paired-end sequencing. Reads of the samples were trimmed for adapters and low-quality bases before alignment with the reference genome (Mouse - mm10) and the annotated transcripts using STAR (36). Differential expression was analyzed using HTSeq and Deseq2 packages (37). The top 50 most differentially expressed genes were selected for hierarchical cluster analysis and visualized using heatmap.

### Statistical Analysis

All data are expressed as the mean ± SD. Differences between groups were evaluated using an unpaired Student’s *t* test. *p* values were considered to be statistically significant when less than 0.05.

## Results

### Increased Th9 differentiation of Bcl-3-deficient CD4^+^ T cells

We have previously demonstrated that Bcl-3 is required for both naïve T cell transfer-induced colitis and experimental autoimmune encephalitis (8). To study how Bcl-3 regulates gene expression in T helper cells, we performed bulk RNA sequencing using wild type (WT) and Bcl-3-deficient (KO) naïve mouse splenic CD4^+^ T cells cultured under Th17 (IL-6 and TGFβ), Th17^+^ (IL-6, TGFβ, IL-21, IL-23 and IL-1β), and Th17^+^ without TGFβ (IL-6, IL-21, IL-23 and IL-1β) conditions. The results showed that *IL-9* was the most highly differentially expressed gene under these conditions (Fig. S1), which suggested that Bcl-3 was involved in regulating IL-9 expression.

Therefore, we explored the role of Bcl-3 in Th9 differentiation. First, we isolated naïve splenic WT and Bcl-3 KO CD4^+^ T cells and cultured them under Th9 conditions for 3 days. FACS analysis consistently showed that the percentage of IL-9^+^ cells increased from ∼10 percent to above 20 percent when Bcl-3 was absent (Fig. 1A, 1B). Consistent with this, real-time PCR analysis showed a two-fold increase of IL-9 mRNA in Bcl-3-deficient Th9 cells compared to WT Th9 cells (Fig. 1C). Interestingly, Bcl-3 KO cells also exhibited higher IL-9 mRNA levels in CD4^+^ T cells cultured under other T helper cell differentiation conditions, even though IL-9 protein was barely detectable in these cells (Fig. S2).

**Figure 1.**
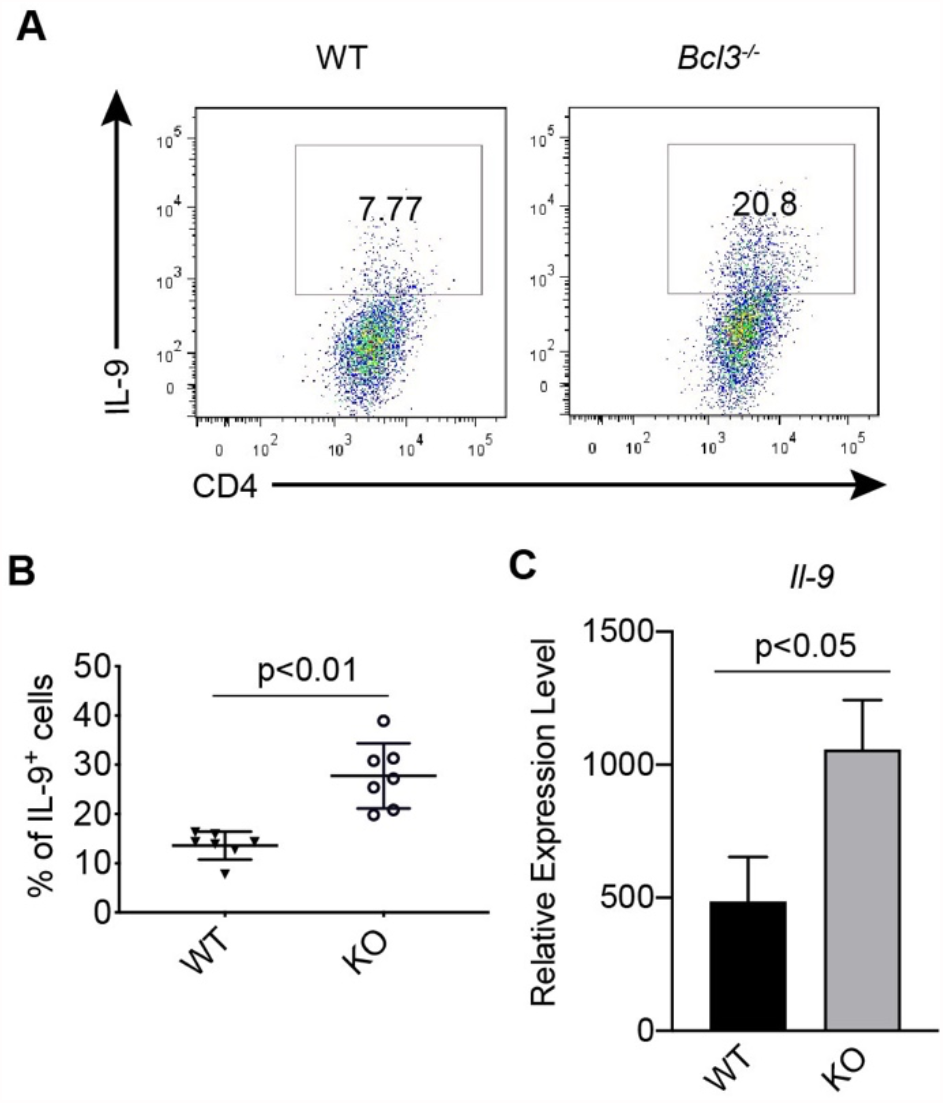
Bcl-3 inhibits Th9 differentiation. (A) Representative flow cytometric analyses of Wild-type (WT) and Bcl-3-deficient (KO) naïve CD4^+^ T cells cultured under Th9 condition for 3 days. (B) Summary of flow cytometric analyses from (A) (n = 7 from two experiments). (C) Real-time PCR analysis of *Il-9* mRNA levels of WT and Bcl-3 KO Th9 cells (n = 3). The data are representative of three independent experiments. Data represent means ± SD.

Next, we isolated naïve WT CD4^+^ T cells from CD45.1 mice and mixed with Bcl-3 KO cells from congenic CD45.2 mice. The mixed cells were then co-cultured under Th9 condition for 3 days and analyzed by FACS. There was still a dramatic increase of Th9 differentiation apparent specifically in Bcl-3 KO cells compared to WT cells in the culture (Fig. 2), which demonstrated that Bcl-3 inhibited Th9 differentiation in a cell-autonomous manner.

**Figure 2.**
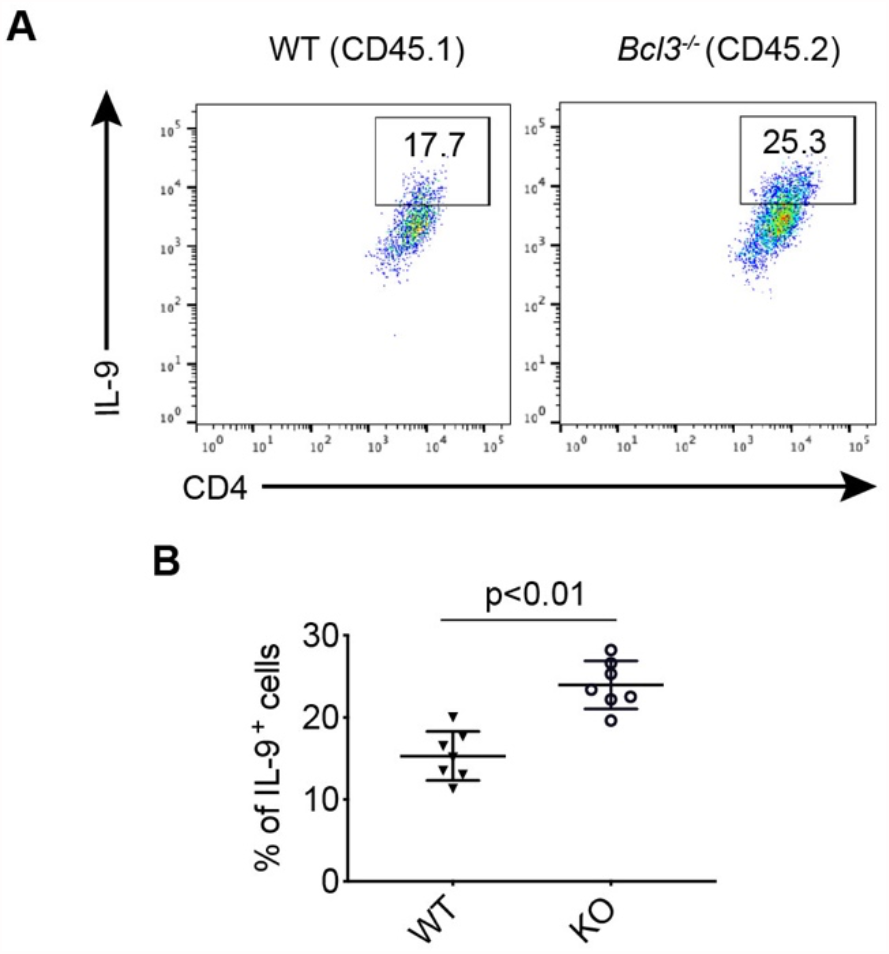
Bcl-3 inhibits Th9 differentiation in a cell-automatous manner. WT (CD45.1) and Bcl-3 KO (CD45.2) naïve CD4^+^ T cells were mixed and cultured under Th9 condition for 3 days. (A) Representative flow cytometric analyses of co-cultured WT and KO cells. (B) Summary of flow cytometric analyses as shown in (A) (n = 7 from three experiments). Data represent means ± SD.

Conversely, we checked whether overexpression of Bcl-3 in T cells might have a reciprocal effect on Th9 cell differentiation. Mice overexpressing Bcl-3 in T cells have been described previously (8). Naïve CD4^+^ T cells from Bcl-3-overexpressing mice and WT littermates were isolated and cultured under Th9 conditions for 3 days. As expected, there was a dramatic decrease of Th9 differentiation in Bcl-3-overexpressing cells (Fig. 3A, 3B). Real-time PCR analysis revealed that Bcl-3 overexpression greatly suppressed IL-9 mRNA expression under Th9 conditions (Fig. 3C).

**Figure 3.**
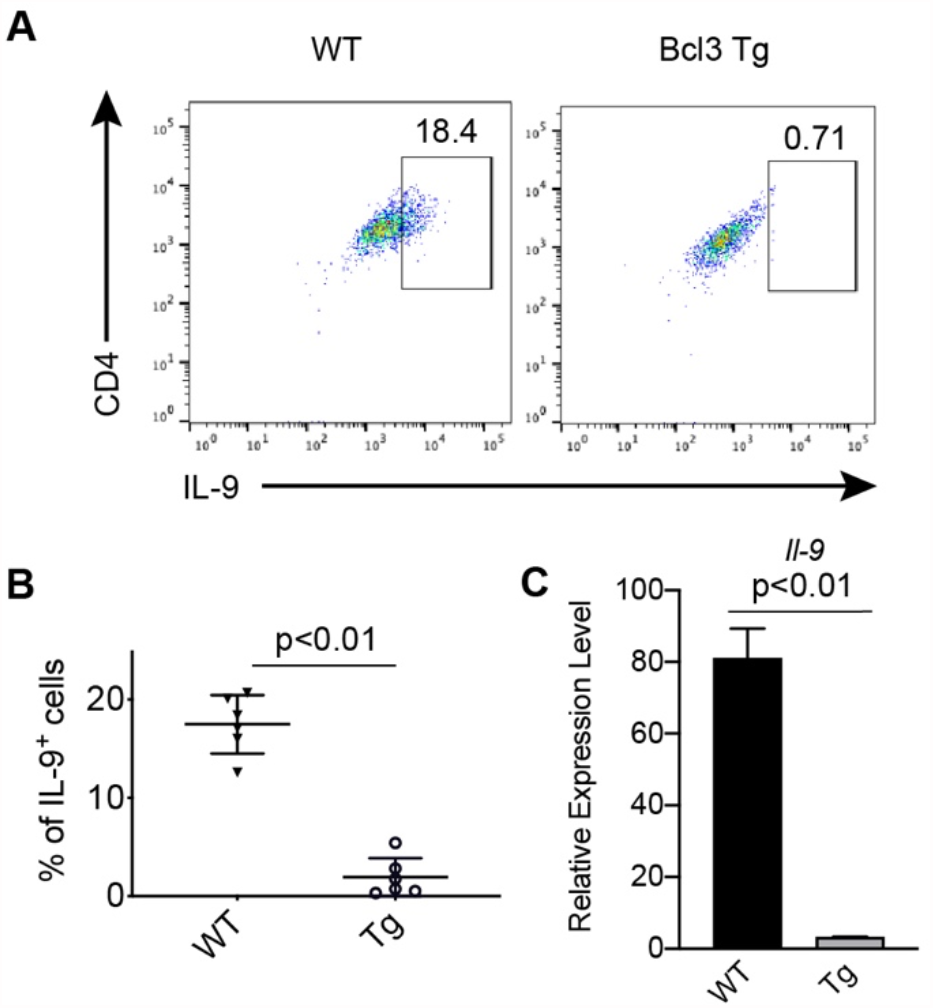
Decreased Th9 differentiation of Bcl-3-overexpressing CD4^+^ T cells. (A) Representative flow cytometric analyses of WT and Bcl-3-overexpressing (Tg) CD4^+^ T cells cultured under Th9 conditions for 3 days. (B) Summary of flow cytometric analyses from (A) (n = 6 from two experiments). (C) Real-time PCR analysis of *Il-9* mRNA levels of WT and Bcl-3-overexpressing Th9 cells. Data are presented as the mean ± SD of 3 replicates from a single experiment representative of 3 independent experiments.

### NF-κB activity is required for increased Th9 differentiation of Bcl-3 KO CD4^+^ T cells

Since Bcl-3 is an atypical IκB protein, we tested whether NF-κB activity was required for increased Th9 differentiation of Bcl-3 KO CD4^+^ T cells. Naive WT and KO CD4^+^ T cells were isolated and cultured under Th9 conditions with different NF-κB inhibitors (Bay 11-7082, IKK Inhibitor VII and MLN-120b) or DMSO as a vehicle control. All three inhibitors greatly suppressed Th9 differentiation, even though they have different targets (Fig. 4). FACS analysis showed a greater suppressive effect of NF-κB inhibitors on Bcl-3 KO cells than on WT cells. MLN-120b exhibited the strongest effect of the three inhibitors, reducing the percentage of Bcl-3 KO IL-9^+^ cells almost to the level of WT. Thus, enhanced Th9 differentiation of Bcl-3 KO cells is strongly dependent on NF-κB signaling.

**Figure 4.**
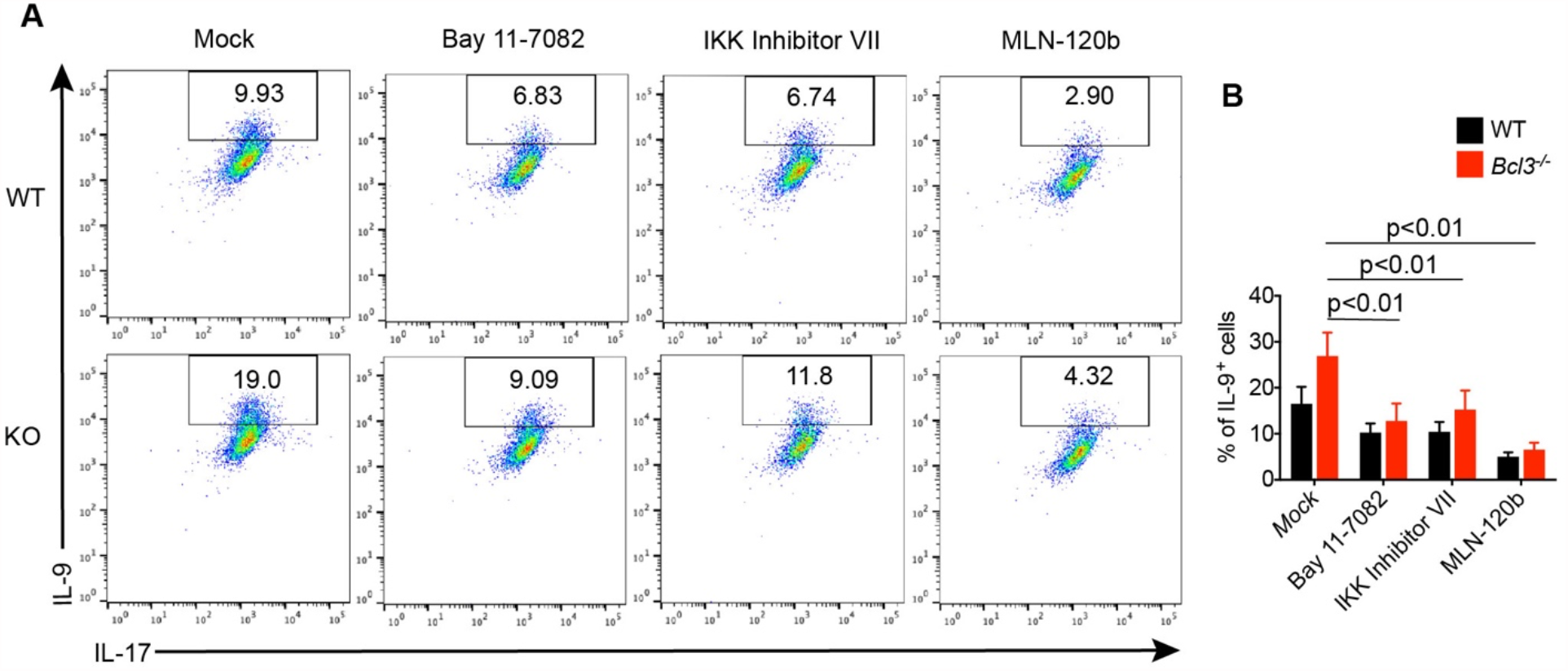
NF-κB activity is required for Th9 differentiation. (A) Representative flow cytometric analyses of WT and Bcl-3 KO CD4^+^ T cells cultured under Th9 conditions with Bay 11-7082 (200 μM), IKK inhibitor VII (400 μM), MLN-120b (5 mM) or DMSO (Mock) for 3 days. (B) Summary data of flow cytometric analyses from (A) (n = 6 from two experiments).

### Bcl-3 suppressed Th9 differentiation by regulating glutamine utilization

To further understand the molecular mechanisms by which Bcl-3 inhibits Th9 differentiation, we performed gene expression assays with WT and Bcl-3 KO Th9 cells. Within 72 h of Th9 stimulation, the expression level of Bcl-3 mRNA first increased and reached a peak at 16 h during Th9 stimulation (Fig. S3A). The early induction of Bcl-3 mRNA provided further evidence that Bcl-3 was involved in Th9 differentiation *in vitro*. We did not detect a significant difference in mRNA levels of different Th9 signature transcription factors (Fig. S3A, S3B).

However, we observed a dramatic reduction of *Slc1a5* mRNA level in Bcl-3 KO cells both before and after Th9 stimulation (Fig. 5A). Slc1a5 is a glutamine transporter and plays an important role in T cell differentiation. To test whether Bcl-3 suppression of Th9 differentiation depends on glutamine utilization, we cultured WT and Bcl-3 KO cells under Th9 conditions with different concentrations of glutamine. FACS analysis showed that the difference between WT and KO cells in the frequency of Th9 cells in the populations decreased when glutamine levels were higher than 8 mM, which supported the idea that Bcl-3 may inhibit Th9 differentiation by controlling glutamine utilization (Fig. 5B, 5C).

**Figure 5.**
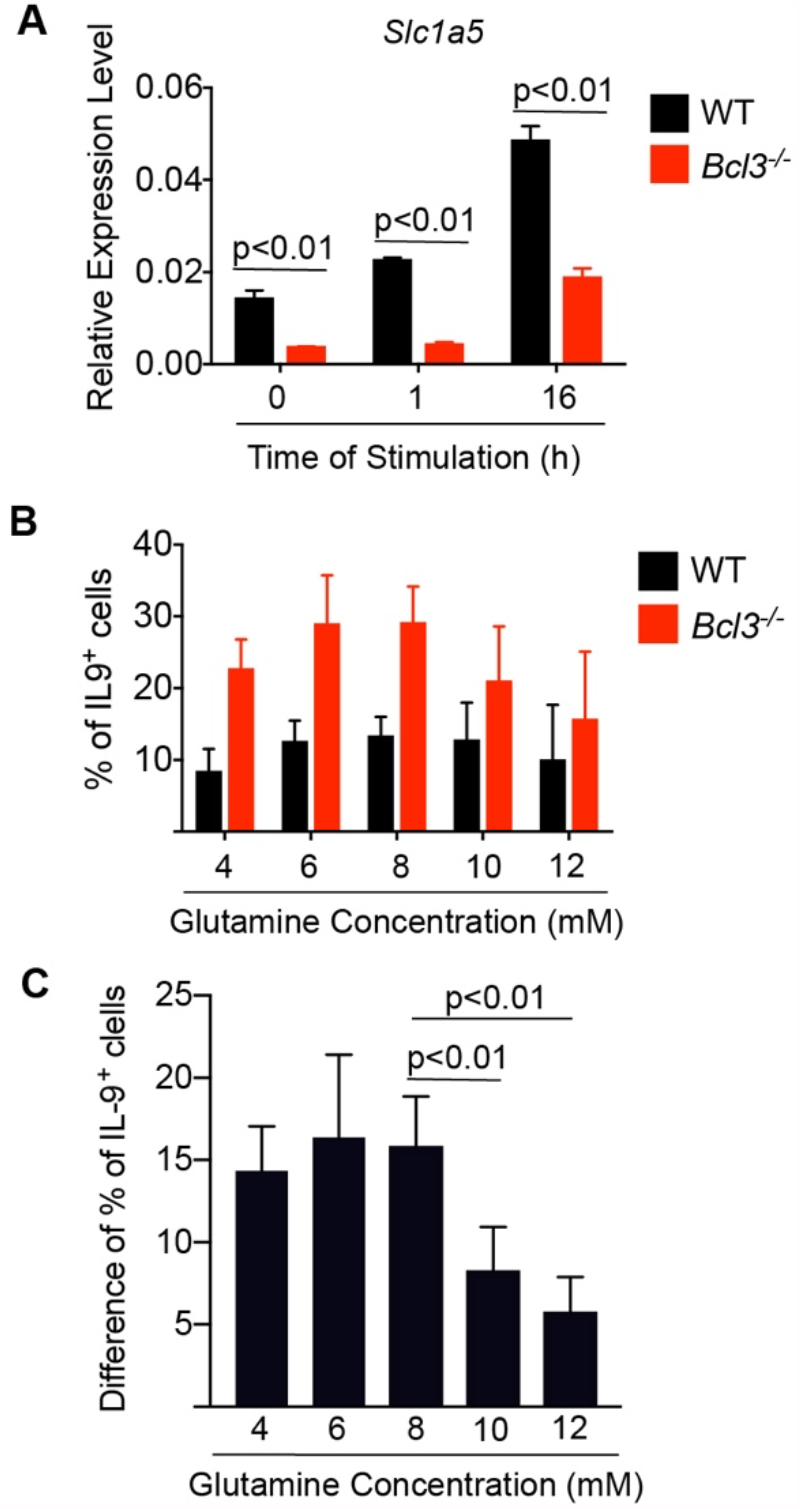
Bcl-3 suppresses Th9 differentiation by regulating glutamine utilization. (A) Real-time PCR analysis of *Slc1a5* mRNA levels of WT and Bcl-3 KO T cells cultured under Th9 conditions. (n = 3 replicates per condition; the experiment was repeated three times with similar results). (B) Summary data of the percentage of IL-9^+^ WT and Bcl-3 KO T cells cultured under Th9 conditions with different concentrations of glutamine (n = 6 from two independent experiments). (C) Difference of the percentage of IL-9^+^ cells between WT and Bcl-3 KO T cells as shown in (B) (n = 6 from two independent experiments). Data represent means ± SD.

### Bcl-3 suppressed lung inflammation in a Th9 transfer model of OVA-induced allergic airway inflammation

We next tested whether the degree of inhibition of Th9 differentiation by Bcl-3 was enough to affect Th9-driven immunopathology *in vivo*. For that, we assessed WT and KO Th9 cells generated in vitro in a Th9 transfer model of OVA-induced allergic airway inflammation. *In vitro* differentiated WT and KO Th9 OTII cells (both CD45.2) were adoptively transferred into WT (CD45.1) mice. The recipients were then challenged intranasally with OVA and IL-25 every 4 days for 24 days and sacrificed on day 25 after the last treatment (Fig. 6A). As expected, intracellular staining of CD45.2 OTII cells showed an increased percentage of IL-9^+^ cells in the lung and mediastinal lymph node of the KO cell recipients (Fig. 6B). More importantly, Bcl-3 KO cell recipients exhibited increased immune cell infiltration of the lung (Fig. 6C, 6D), which was consistent with the notion that Bcl-3 KO Th9 cells are more pathogenic *in vivo*.

**Figure 6.**
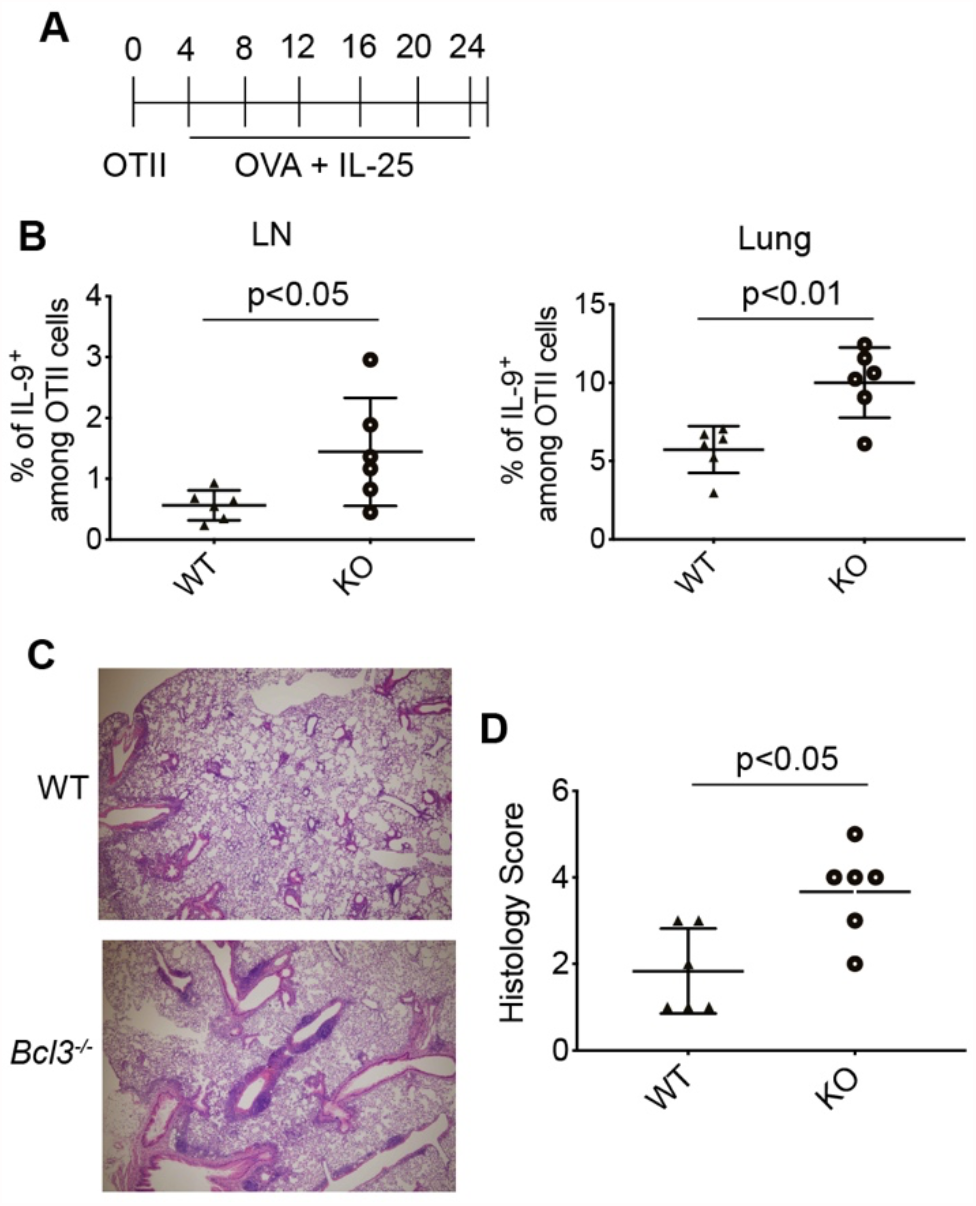
Bcl-3 inhibits immunopathology in a Th9 transfer model of OVA-induced allergic airway inflammation. (A) Timeline of Th9 transfer model of OVA-induced allergic airway inflammation: WT and Bcl-3 KO OT-II cells were cultured under Th9 conditions *in vitro* and adoptively transferred into CD45.1 mice. The recipients were treated with Ovalbumin and IL-25 every 4 days for 24 days and euthanized one day after the last treatment. (B) Summary of flow cytometric analyses of IL-9^+^ cells of lung and mediastinal lymph nodes from the recipients (n = 6 from two experiments). (C) Representative H&E-stained sections of lungs. (D) Summary of histology score of lung sections from (C) (n = 6 from two experiments). Data represent means ± SD.

## Discussion

In the present study, we provide evidence that the I*k*B family member Bcl-3 negatively regulates IL-9 expression at the RNA level resulting in reduced Th9 cell differentiation of naïve CD4^+^ T cells. The magnitude of the effect is sufficient to diminish Th9-dependent OVA-induced allergic immunopathology in the lung in an adoptive transfer model. The mechanism may involve in part reduced accessibility to glutamine due to downregulation of the glutamine transporter *Slc1a5*.

Our hypothesis that Bcl-3 regulates Th9 differentiation originated with results from an unsupervised transcriptomic analysis of Th17 differentiation of Bcl-3 knockout CD4^+^ T cells *in vitro* that was conducted as part of a mechanistic investigation of our previously published finding that Bcl-3 is required for both naïve T cell transfer-induced colitis and experimental autoimmune encephalitis (8). Although *IL-9* was the most highly differentially expressed gene and was upregulated in Bcl-3 knockout Th17 cells, still IL-9 mRNA was expressed at very low levels and IL-9 protein was barely detectable in both wild type and KO Th17 cells. The same pattern of Bcl-3-dependent IL-9 expression was observed in CD4^+^ T cells cultured under Th1 and Treg conditions *in vitro*. Thus, although IL-9 is a minor component of Th1, Th17 and Treg cells and a major and defining component of Th9 cells, it is negatively regulated by Bcl-3 in all four cell types. In contrast, we have shown previously that Th1, Th17 and Treg differentiation is not affected by Bcl-3 deficiency (8, 33).

Our results suggest a general mechanism of Bcl-3-dependent regulation of *IL-9* and perhaps other genes, regardless of the cytokine environment. The mechanism appears to involve the canonical NF-*κ*B pathway, since three different pharmacological inhibitors of this pathway all markedly inhibited enhanced Th9 differentiation in Bcl-3 KO cells. This is consistent with the known ability of Bcl-3 to bind to NF-*κ*B homodimers of p50 and p52, and in this way to regulate NF-κB activity on DNA. The non-canonical NF-κB pathway might also contribute to the Bcl-3 effect and will require further investigation. In this regard, it has previously been reported that OX40-mediated signaling promotes Th9 differentiation via enhanced non-canonical NF-κB activity. OX40 also activates the canonical NF-κB pathway but canonical NF-κB activity is not required for OX40L-induced Th9 differentiation (38). This discrepancy may be due to compensation by non-canonical NF-κB activity when the canonical NF-κB pathway is inhibited during OX40L stimulation. Therefore, a prediction from our results for future investigation is that OX40-mediated signaling may reverse the inhibitory effect of Bcl-3 on Th9 differentiation.

Further studies on Bcl-3 binding sequences and interacting proteins are required to better understand the mechanism of how Bcl-3 regulates Th9 differentiation. To date, progress in this regard has been hindered by the lack of suitable antibody reagents for detection of Bcl-3. Additional mechanisms mediating Bcl-3 regulation of Th9 differentiation that are independent of NF-*κ*B might also be in play and should be considered, including its known association with HDACs (39). Our unsupervised transcriptomic analysis of wild type versus Bcl-3-deficient Th9 cells identified the glutamine transporter *Slc1a5* as a highly differentially expressed gene that was down-regulated in different types of Bcl-3-deficient Th cells. The ability of glutamine excess to reverse Bcl-3-dependent regulation of Th9 differentiation suggests a novel metabolic mechanism mediating Bcl-3 action. There is no report on the role of glutamine in Th9 differentiation so far. However, it has been reported that glutamine promotes Th1 but restrains Treg differentiation (40). And the glutamine transporter *Slc1a5* has been shown to be critical for pathogenic Th1 and Th17 cells (41). Our results provided evidence that glutamine was also involved in the regulation of Th9 differentiation. Further studies are needed to define the metabolic intermediates involved in this process.

Our results have potential clinical implications in that Th9 cells have been reported to be effective in adoptive cell transfer therapy of cancer, and more effective than Th1 or Th17 cells (27). Since Bcl-3 inhibits Th9 differentiation, it is a potential molecular target to further enhance the efficacy of Th9 cell cancer therapy. In this regard, one way of leveraging our discovery of Bcl-3 regulation of Th9 differentiation may be to simply manipulate the glutamine concentration during the *ex vivo* differentiation process.

## Acknowledgements

We greatly appreciate the constructive inputs provided by all members of the Siebenlist laboratory. This research was supported by the Division of Intramural Research, NIAID, NIH.

