## Supplemental Data for "Bcl-3 suppresses Th9 differentiation by regulating glutamine utilization"

Figure S1.

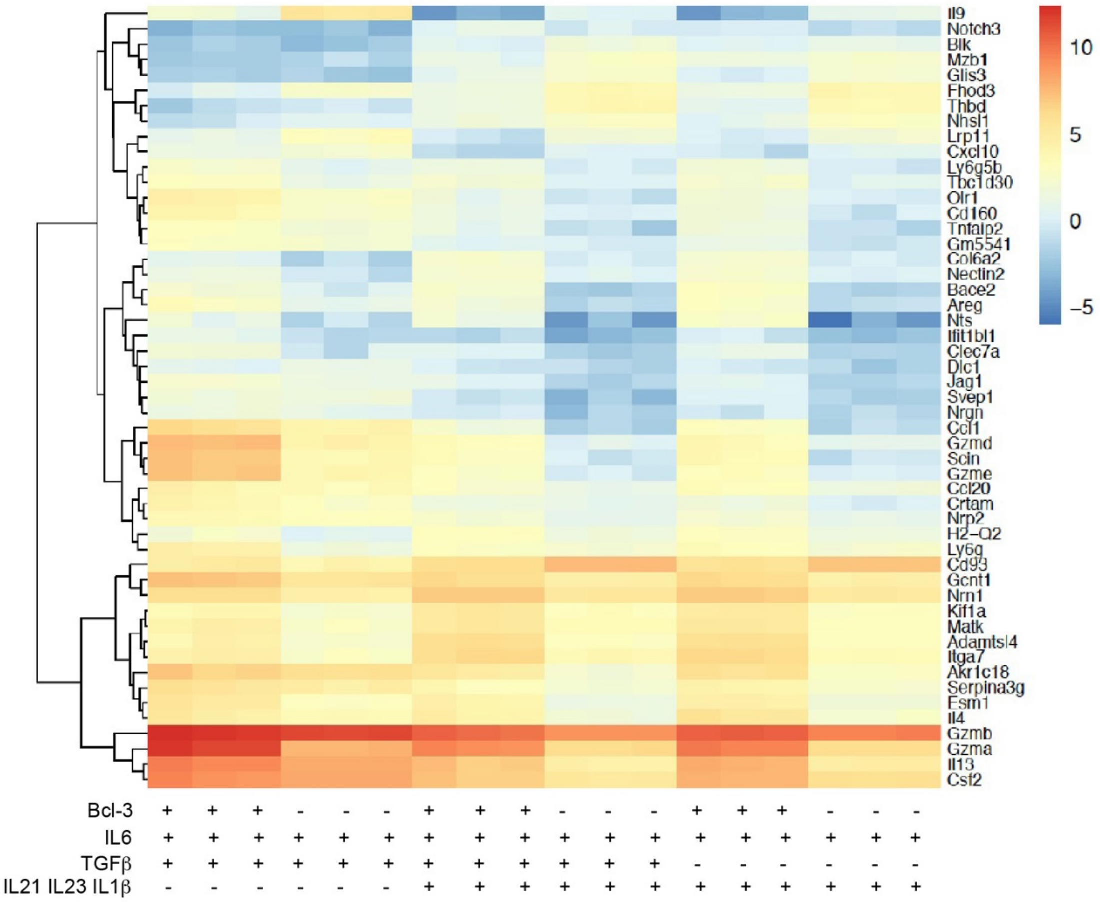


Figure S1. Top 50 most differentially expressed genes between WT and Bcl-3 KO cells cultured under different Th17 conditions.

Figure S2.

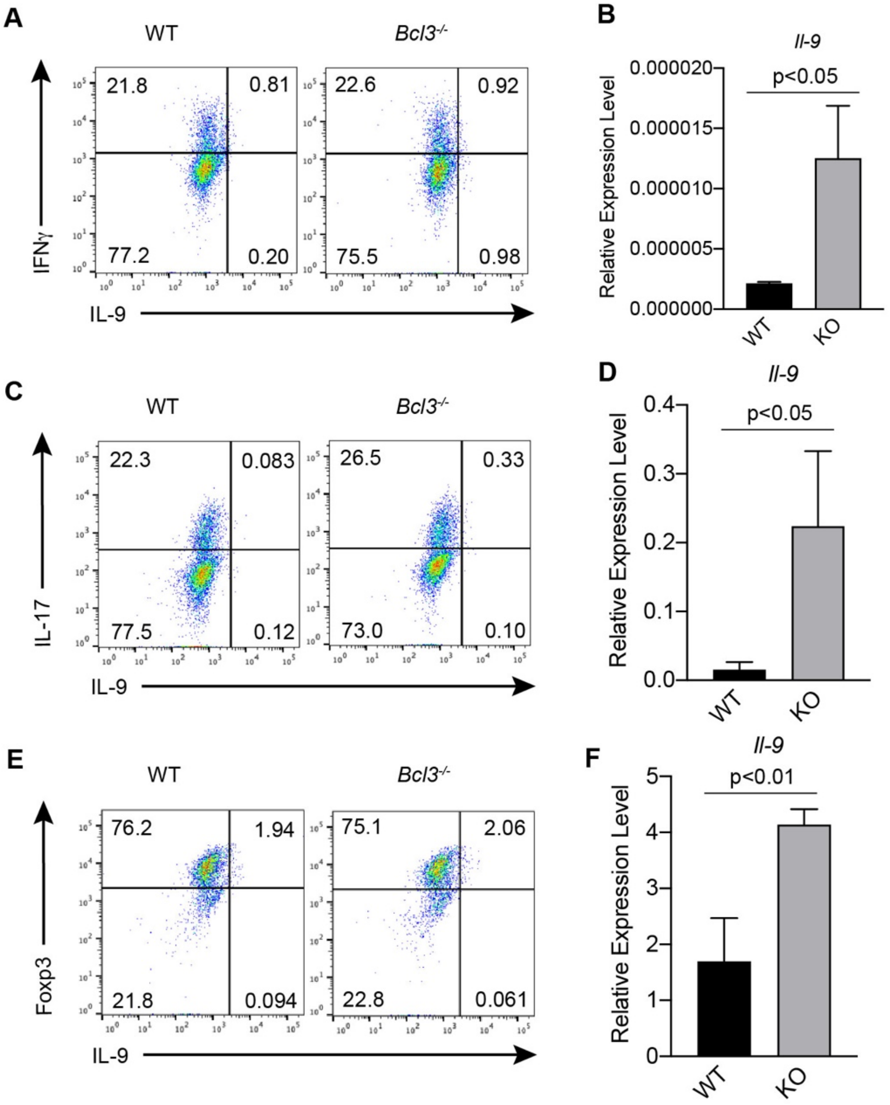


Figure S2. Increased *Il-9* mRNA levels in different Bcl-3 KO T helper cells. (A, C and E) Representative flow cytometric analyses of WT and Bcl-3 KO Th1, Th17 and Treg cells. (B, D and F) Real-time PCR analysis of *Il-9* mRNA levels of WT and Bcl-3 KO Th1, Th17 and Treg cells (n = 3, these experiments were repeated 3 times).

Figure S3.

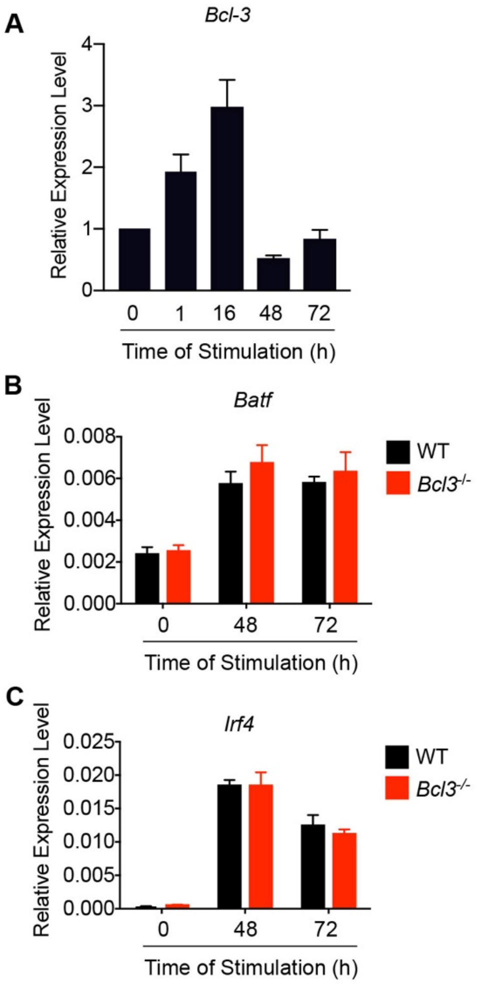


Figure S3. No significant difference in mRNA levels of Th9 signature transcription factors between WT and Bcl-3 KO Th9 cells. (A) Real-time PCR analysis of *Bcl-3* expression in WT cells during Th9 differentiation. (B and C) Real-time PCR analysis of *Batf* and *Irf4* expression in WT and Bcl-3 KO Th9 cells.
